# A selection of Golden Gate vectors to simplify recombinant protein production in *Escherichia coli*

**DOI:** 10.1101/2024.02.13.579886

**Authors:** Michael Fairhead, Lizbe Koekemoer, Eleanor Williams, Frank von Delft

**Affiliations:** Centre for Medicines Discovery, Nuffield Department of Medicine, NDM Research Builiding, University of Oxford, Old Road Campus, Oxford, OX3 7FZ

## Abstract

In this work multiple plasmids have been created to allow the simple Golden Gate cloning of a target gene for recombinant protein production in *Escherichia coli*. To simplify as much as possible the generation of different target gene vector combinations, the 22 plasmids contain the same Golden Gate cloning sites (*Bsa*I), antibiotic resistance (kanamycin) and promoter (T7) for expression in the standard protein production strain of *E. coli* BL21[DE3]. The plasmid set includes commonly used tags for purification and assays (his, twin-strep and avi tag) as well as fusion protein partners that may aid target protein solubility and yield, SUMO, MBP, GST and sfGFP. Also included are plasmids with secretion peptide signals for transport of the target protein to the *E. coli* periplasm *via* various pathways (SEC, SRP, TatA).

We have evaluated the 4 of the vectors using a test super folder GFP insert and found that using the Golden Gate process allows cloning efficiencies of greater than 90% to be routinely obtained. Vectors were further evaluated by expressing and purifying the target insert. The plasmid vector set described herein should prove useful to any investigator who has to routinely evaluate numerous protein expression constructs and are freely available through Addgene.

## Introduction

Methods for the cloning of target inserts into vectors suitable for recombinant protein production are many and varied, from the “classical” restriction enzyme digestion and ligation method first described in the1980’s (1) to more recent PCR based methods such as restriction free cloning (2) and Gibson assembly (3). Many of these systems have advantages and disadvantages with some such as LIC requiring specific treatments of the vector and insert (4) and others requiring very long primers and multiple PCR reactions(2).

The Golden Gate Cloning method (5) enables the simple cloning of a target DNA insert into multiple vector types for recombinant protein production as has been shown by Rohweder *et al*. (6) and simply requires the mixture of a compatible insert and vector pair in a single reaction for cloning. The method relies on the use of a type IIs restriction enzyme (e.g. *Bsa*I) which can cut down- or up-stream of their recognition sequence (depending on the orientation in target DNA) and subsequent ligation of the cut DNA by T4 DNA ligase, see Figure 1.

**Figure 1.**
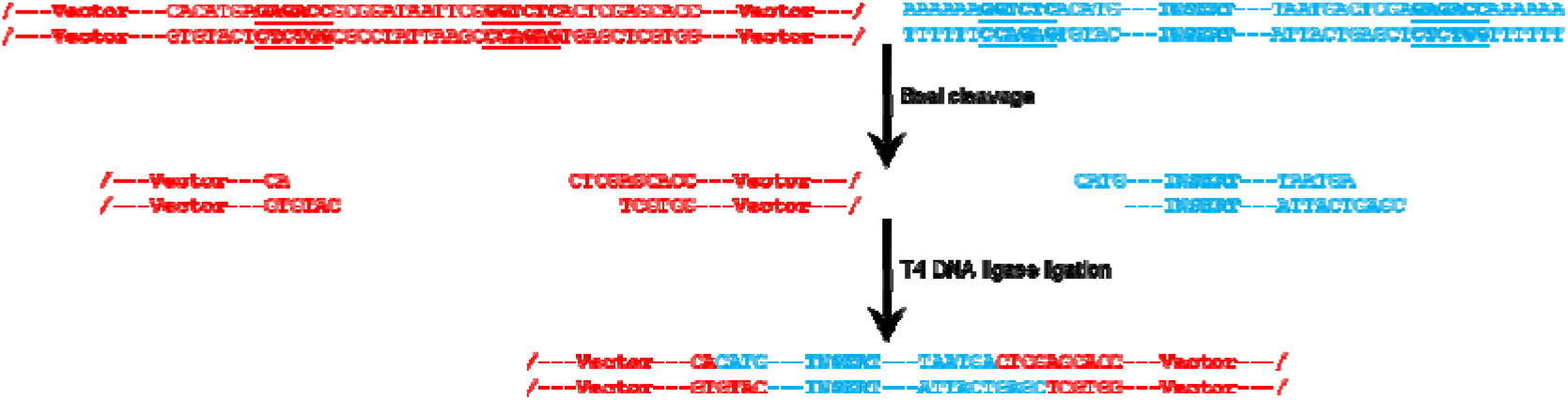
Principle of Golden Gate cloning. Both the vector and insert are cut with the type IIs *Bsa*I restriction enzyme and the compatible DNA overhangs are subsequently ligated together by the action of T4 DNA ligase.

This elegantly simple process allows the seamless ligation of many vector insert combinations leaving minimal or no multiple cloning site artefacts such as unwanted tags and linkers, which can be hard to avoid generating when doing standard restriction cloning.

The vector set described in this work (see Table 1.) covers many of the tags commonly desired by experimentalists such as cleavable poly histidine (7) and twin strep tags (8) for purifications, solubilizing fusion proteins such as MBP (9), GST (10) and SUMO (11) and the avi-tag (12) which allows site specific biotinylation of the target protein for assays and biophysics. We have also included a selection of less commonly used tags including thioredoxin (13), nusA (14) and Zbasic (15). Vectors for targeting the recombinant protein to the *E. coli* periplasm via the SEC, SRP and TatA pathways (16) have also been made.

**Table 1.**
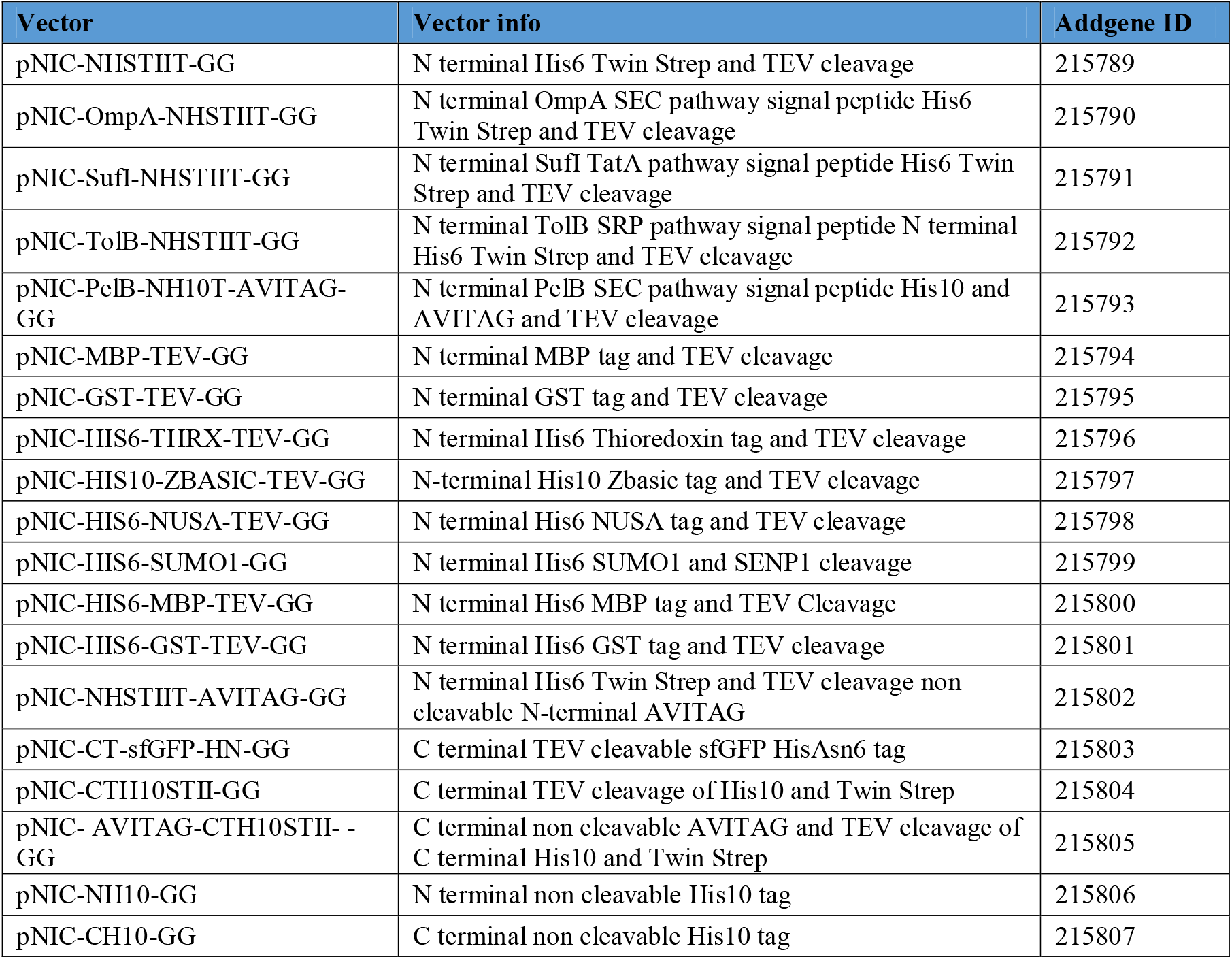
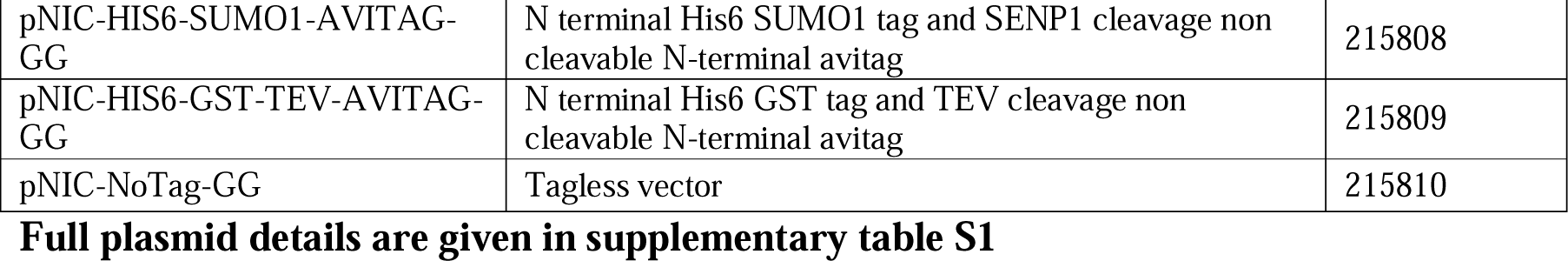
List of golden gate vectors generated in this work.

## Materials & Methods

### Plasmid construction

Plasmids were typically generated either *via* modification of a pre-existing in house vector using mutagenesis or by cloning a synthetic gene insert (gBlock from Integrated DNA Technologies) into the *Nde*I and *Sal*I restriction enzyme sites of pNIC28-*Bsa*4 (17). Plasmids were initially verified *via* Sanger sequencing (Source Bioscience) using T7forward (5’-TAATACGACTCACTATAGGG-3’) and T7reverse (5’-GCTAGTTATTGCTCAGCGG-3’) primers and then fully sequenced using Nanopore sequencing (Source Bioscience). Virtual plasmid vector maps and sequences were constructed using the SnapGene software (Dotmatics). *In-silico* validation of the Golden Gate cloning was also done using SnapGene.

### Primers

The target gene was first amplified with a forward primer containing the 5’ golden gate extension, 5’-AAAAAA<u>GGTCTC</u>AC*ATG*–target-3’, where the *Bsa*I site is underlined and the *ATG* is the start codon of the target gene. The annealing temperature (Tm) of the target section of the primer is in the region of 55-60°C and was calculated using the NEB Tm calculator (https://tmcalculator.neb.com/#!/main). The reverse primer followed a similar design principle but the 5’ extension was, 3’-target-*TAATGA*CTCGA<u>GAGACC</u>AAAAAA-5’, complement 5’-TTTTTTGGTCTCTCGAG*TCATTA*-tegrat-3’. For c-terminal tag vectors the stop codons *TAATGA* were removed from the primer, while the remainder of the sequence was the same.

### Polymerase chain reaction to amplify insert

PCR amplification of the insert was done with Q5 DNA polymerase from NEB. The standard Q5 reaction contained 5 μl of 5x Q5 reaction buffer, 1.25 μl of forward primer (10 μM), 1.25 μl of reverse primer (10 μM), 0.5 μl of dNTPs (10 mM), 1 μl of template DNA (5 ng/μl), nuclease free water to a final volume of 25 μl and finally 0.25 μl of Q5-HF polymerase.

The following protocol was then run on a T100 thermocylcer (BioRad) using a heated lid (105 °C). An initial denaturation step for 30 seconds at 98°C followed by 30 cycles of 10 seconds as 98 °C, 20 seconds of annealing at the primer Tm and 30 seconds of extension per kb of insert at 72°C. This was followed by a 2 minute polishing step at 72°C after which the reaction was held at 12°C until further processing. The fidelity of the PCR reaction was checked using a 1 % agarose gel and then the reaction cleaned up using a PCR clean up spin column kit (Qiagen). Finally the concentration of the purified PCR product was checked on a NanoDrop spectrophotometer (Thermo-Fisher).

If using template plasmid DNA that had the same antibiotic resistance as the destination golden gate vector (kanamycin) a *Dpn*I digest to remove the template post PCR was necessary. To do this 2.5 uL of cutsmart buffer was added to 20 uL of purified PCR product and then 1 uL of *Dpn*I (NEB) was added. The reaction was incubated at 37°C for 30 minutes to digest methylated template plasmid DNA, followed by an incubation at 80°C for 20 minutes to inactivate the *Dpn*I enzyme after which the reaction was kept on ice until further processing.

### Golden gate assembly reaction

The golden gate assembly reaction was assembled in a PCR tube by mixing 100 ng of the vector with a 3 fold molar excess of the purified PCR product (calculated using the NEB bio calculator, available at https://nebiocalculator.neb.com/#!/ligation) after which water was added to a final volume of 15 μl followed by 1 μl of 20 mg/mL BSA, 2 μl of 10 x T4 DNA ligase buffer, 1 μl of *Bsa*I-HFv2 and 1 μl of T4 DNA ligase (NEB).

The following protocol was then run on a T100 thermocylcer (BioRad) using a heated lid (105°C). The reaction was first held for 10 minutes at 37°C followed by 5 cycles of 5 minutes at 37°C and 5 minutes at 16°C. This was followed by 5 minutes at 37°C and then 20 minutes at 80°C after which the reaction was held at 12°C until further processing. An aliquot of the reaction (3 μL) was then used to transform 100 μL of home-made CCMB80 (18) MachI *E. coli* (Thermo Fisher) competent cells. After selection on LB-agar kanamycin plates the resulting colonies were first screened by colony PCR using MyTaq DNA polymerase (Bioline) to check for the presence of the right size insert using T7forward and T7reverese primers. Sanger sequencing using the T7forward (5’-TAATACGACTCACTATAGGG-3’)and T7reverse (5’-GCTAGTTATTGCTCAGCGG-3’) primers was then used to check the final cloned insert sequence.

### Test Expression and purification

A 100 mL test expression and purification was performed using the PREPX workflow (19), briefly described below. Typically 100 ng of plasmid was transformed into the *E. coli* protein expression strain BL21(DE3) and a single colony used to inoculate 10 mL of SOC + 50 μg/mL kanamycin. The starter culture was grown overnight at 37°C 250 rpm shaking and the following day 1 mL was used to inoculate 100 mL of AIM-TB (ForMedium) in a 250 mL baffled shake flask supplemented with 50 μg/mL kanamycin and 0.01% Antifoam-204 (Merck). Cells were grown first for 4 h at 37°C and then for a further 20 h at 18°C using 250 rpm shaking. Cells were harvested by centrifugation at 4,000 g and the pellets re-suspended using 35 mL of Base Buffer (10 mM HEPES, 500 mM NaCl, 5 % glycerol, 30 mM imidazole 0.5 mM TCEP, pH 7.5) per 100 mL culture. Triton X-100, lysozyme and benzonase to a final concentration of 1 %, 0.5 mg/mL and 1 μg/mL respectively were then added before freezing at -80°C. The cell lysate mixture was first thawed in a room temperature water bath and then centrifuged at 4800 g for 1 h at 4°C. The supernatant (soluble fraction) was then loaded onto a 1 mL His GraviTrap column (Cytiva) and washed with 3 times with 10 mL of Base Buffer. The tagged-target protein was then eluted using 2.5 mL of Base Buffer + 500 mM imidazole and immediately desalted using a PD-10 column (Cytiva). Where appropriate the tag was then removed by protease digest overnight at 4°C using either 1 mg of TEV per 10 mg of target protein or in the case of SUMO fusions, 1 mg of SENP1 per 100 mg of target. The tag, uncleaved protein and proteases (bearing a non-cleavable his tag) was then removed by passing through a His GraviTrap column. Efficacy of the process and final protein purity was assessed by SDS-PAGE using NuPAGE 4-12 % Bis-Tris Midi Protein Gels (ThermoFisher Scientific).

## Results

The main aim of this work was to create a suite of plasmid vectors to simplify the cloning of a target gene and its subsequent recombinant production in *E. coli*. To achieve this goal 22 plasmid vectors were created encoding the various tags, fusion partners and signal peptides commonly used in *E. coli* recombinant protein production, see Table 1. After cloning all plasmid sequences were verified *via* Nanopore sequencing and validated *in-silico* using the Golden Gate cloning function of SnapGene.

A real-world test of some of the new vectors and the Golden Gate cloning method was done using a super folder GFP (sfGFP) (20) insert. The sfGFP insert was cloned into four different plasmids pNIC-NHSTIIT-GG, pNIC-HIS6-SUMO-GG, pNIC-HIS6-GST-TEV-GG, pNIC-CTH10STII-GG and the success of the Golden Gate cloning workflow (see Methods) was assessed by colony PCR. T7Forward and T7reverse primers were used to check for the absence or presence of the sfGFP insert in the clones generated after transforming competent MachI *E. coli* with an aliquot of the golden gate reaction mixture, Table 2 and supplementary figures 1-4. As judged by the colony PCR outcomes for the four vectors tested successful cloning was achieved around 90 % of the time with some vector to vector variation evident. One positive clone for each of the four vector insert combinations was subsequently sent for Sanger sequencing using the T7forward and T7reverse primers. Sequencing confirmed the presence of the sfGFP insert and its plasmid encoded tag (supplementary sequences 1-4).

**Table 2.** Efficacy of the Golden Gate cloning workflow as judged by colony PCR outcome for sfGFP clones.

| **VECTOR** | **Positive colony PCRs** |
| --- | --- |
| pNIC-NHSTIIT-GG | 19/23 (83 %) |
| pNIC-HIS6-SUMO-GG | 22/23 (96 %) |
| pNIC-HIS6-GST-TEV-GG | 21/23 (91 %) |
| pNIC-CTH10STII-GG | 22/23 (96 %) |
|  | **Total 84/92 (91 %)** |

A small scale test expression and protein purification was then done for each of the four sfGFP constructs using the PREPX workflow (19) to ensure the encoded plasmid tags behaved as expected for the purposes of recombinant protein production. In Figure 2. is shown the outcome of this experiment for the pNIC-NHStIIT-sfGFP construct, evaluated using SDS-PAGE. It can be seen that the tagged protein (32.5 kDa) is overexpressed in whole cells (Figure 2, lane WC) and that the protein expressed is predominantly in the soluble fraction (Figure 2, lane SF). The soluble expressed protein can be captured on an IMAC His Gravitrap column using the plasmid encoded His tag sequence (Figure 2, lane IM). The His twin strep tag can then be cleaved off the protein specifically using TEV protease (21) and the tag and any un-cleaved protein removed using a reverse IMAC process (Figure 2, lane RV) yielding relatively pure untagged protein (26.9 kDa). The other three sfGFP plasmids generated (pNIC-HIS6-SUMO-GG, pNIC-HIS6-GST-TEV-GG, pNIC-CTH10STII-GG) were also found to perform as expected, see supplementary Figures 5, 6 and 7 as judged by SDS-PAGE. In summary all four sfGFP tagged constructs created were well expressed and exhibited the expected properties in terms of molecular weight, his tag purification and protease based tag removal.

**Figure 2.**
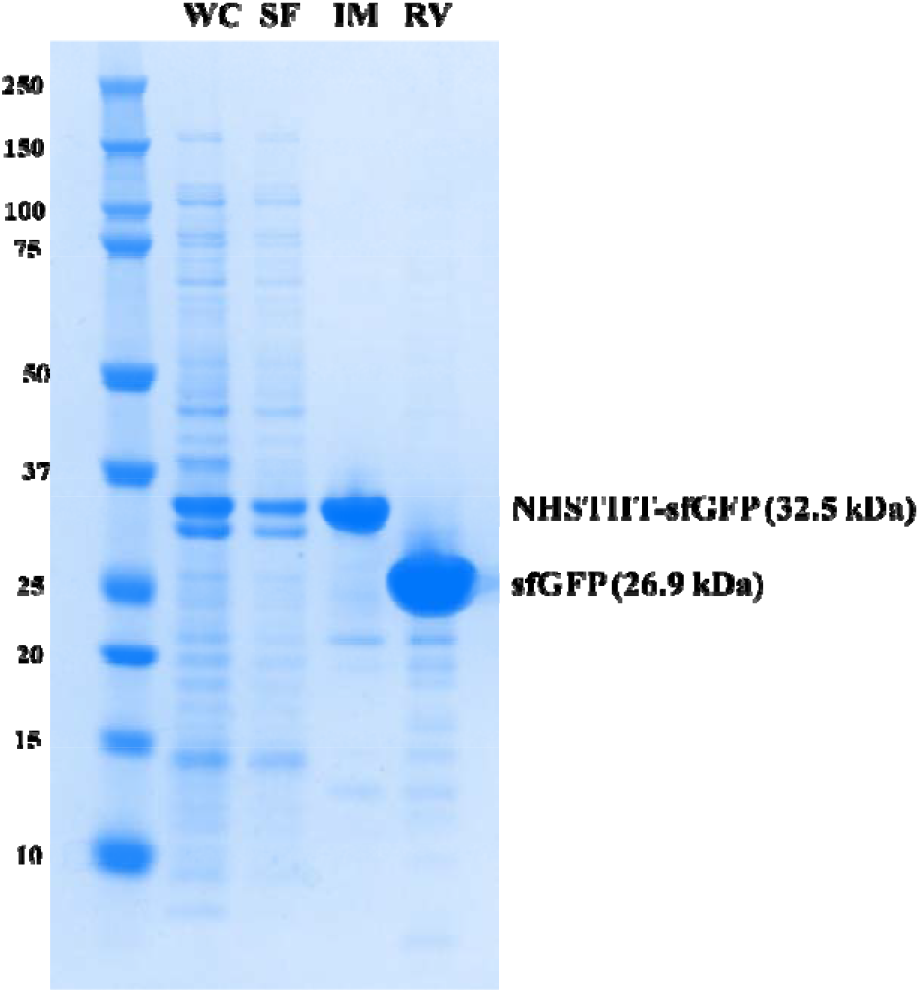
SDS-PAGE of the pNIC-NHSTIIT-sfGFP construct protein purification, where WC = whole cell lysate, SF = soluble fraction obtained after centrifugation, IM = 500 mM imidazole His Gravitrap eluate, RV = His Gravitrap flow-through after tag removal by protease cleavage.

## Discussion and Conclusions

In this work a total of 22 plasmid vectors for recombinant protein production in *E. coli* were created. The vectors have been designed in such a way as to simplify the process of evaluating the effect of different solubility and purification tag combinations on recombinant protein yield and purity. As all plasmids contain the same cloning site, the simultaneous evaluation of multiple target insert vector combinations is relatively straightforward, although it is necessary for inserts going into N-terminal tagged vectors to encode a stop codon. One obvious major limitation would be the presence of a *Bsa*I site in the insert gene sequence. This situation can be avoided by ordering codon optimised synthetic gene inserts without any internal *Bsa*I from suppliers such as TWIST or IDT. As we and others have found the Golden Gate method is particularly well suited to the direct cloning of synthetic gene fragments (22). For the system described in this work provided the synthetic genes are ordered with the above mentioned golden gate extensions (5’-AAAAAA<u>GGTCTC</u>AC*ATG*-target-*TAATGA-* CTCGA<u>GAGACC</u>AAAAAA-3’) the received insert(s) can then be used directly in a Golden Gate reaction with the vector of choice.

The full details of the created plasmids and their sequences are given in supplementary Table S1, in the same file is also a vector insert cloning spreadsheet, Table S2. The spreadsheet allows the simple generation of the DNA and amino acid sequences produced when the target insert is combined with one of the various vectors described in this work. The sheet also generates the new constructs predicted molecular weight and extinction co-efficient before and after tag removal.

In conclusion we believe the set or plasmids described in this work should be useful to anyone wishing a quick and straightforward method to evaluate the feasibility of producing a target protein in *E. coli*.

## Supporting information

Manuscript Figures

Vector properties and cloning sheet

## Acknowledgements

MF was supported by BBSRC grant no. BB/T003677/1

**Supplementary sequence 1**. Amino acid sequence of the NHSTIIT-sfGFP construct. Calculated MW of NHSTIIT-sfGFP is 32.5 kDa and after TEV cleavage (*) and tag removal, 26.9 kDa.

MHHHHHHSSGASWSHPQFEKGGGSGGGSGGSAWSHPQFEKGSGVDLGTENLYFQ*SMSKGEE LFTGVVPILVELDGDVNGHKFSVRGEGEGDATNGKLTLKFICTTGKLPVPWPTLVTTLTYGVQCFSRYPDHMKRHDFFKSAMPEGYVQERTISFKDDGTYKTRAEVKFEGDTLVNRIELKGIDFKEDGNILGHKLEYNFNSHNVYITADKQKNGIKANFKIRHNVEDGSVQLADHYQQNTPIGDGPVLLPDNHYLSTQSVLSKDPNEKRDHMVLLEFVTAAGITHGMDELYK

**Supplementary sequence 2**. Amino acid sequence of the His6-SUMO-sfGFP construct. Calculated MW of His6-SUMO-sfGFP is 38.8 kDa and after SUMO cleavage (*) and tag removal, 26.8 kDa.

MHHHHHHGSGDQEAKPSTEDLGDKKEGEYIKLKVIGQDSSEIHFKVKMTTHLKKLKESYCQR QGVPMNSLRFLFEGQRIADNHTPKELGMEEEDVIEVYQEQTGG*MSKGEELFTGVVPILVEL DGDVNGHKFSVRGEGEGDATNGKLTLKFICTTGKLPVPWPTLVTTLTYGVQCFSRYPDHMKRHDFFKSAMPEGYVQERTISFKDDGTYKTRAEVKFEGDTLVNRIELKGIDFKEDGNILGHKLEYNFNSHNVYITADKQKNGIKANFKIRHNVEDGSVQLADHYQQNTPIGDGPVLLPDNHYLSTQSVLSKDPNEKRDHMVLLEFVTAAGITHGMDELYK

**Supplementary sequence 3**. Amino acid sequence of the His6-GST-sfGFP construct. Calculated MW of His6-GST-sfGFP is 54.6 kDa and after TEV cleavage (*) and tag removal, 26.9 kDa

MHHHHHHSSMSPILGYWKIKGLVQPTRLLLEYLEEKYEEHLYERDEGDKWRNKKFELGLEFPNLPYYIDGDVKLTQSMAIIRYIADKHNMLGGCPKERAEISMLEGAVLDIRYGVSRIAYSKDFETLKVDFLSKLPEMLKMFEDRLCHKTYLNGDHVTHPDFMLYDALDVVLYMDPMCLDAFPKLVCFKKRIEAIPQIDKYLKSSKYIAWPLQGWQATFGGGDHPPKSGGGSENLYFQ**\***SMSKGEELF TGVVPILVELDGDVNGHKFSVRGEGEGDATNGKLTLKFICTTGKLPVPWPTLVTTLTYGVQCFSRYPDHMKRHDFFKSAMPEGYVQERTISFKDDGTYKTRAEVKFEGDTLVNRIELKGIDFKEDGNILGHKLEYNFNSHNVYITADKQKNGIKANFKIRHNVEDGSVQLADHYQQNTPIGDGPVLLPDNHYLSTQSVLSKDPNEKRDHMVLLEFVTAAGITHGMDELYK

**Supplementary sequence 4**. Amino acid sequence of the sfGFP-CTH10STII construct. Calculated MW of sfGFP-CTH10STII is 32.2 kDa and after TEV cleavage (*) and tag removal, 26.7 kDa

MSKGEELFTGVVPILVELDGDVNGHKFSVRGEGEGDATNGKLTLKFICTTGKLPVPWPTLVT TLTYGVQCFSRYPDHMKRHDFFKSAMPEGYVQERTISFKDDGTYKTRAEVKFEGDTLVNRIE LKGIDFKEDGNILGHKLEYNFNSHNVYITADKQKNGIKANFKIRHNVEDGSVQLADHYQQNT PIGDGPVLLPDNHYLSTQSVLSKDPNEKRDHMVLLEFVTAAGITHGMDELYKLENLYFQ*SH HHHHHHHHHGSGWSHPQFEKGGGSGGGSGGSAWSHPQFEK

## Supplementary Figures

**Supplementary Figure 1.**
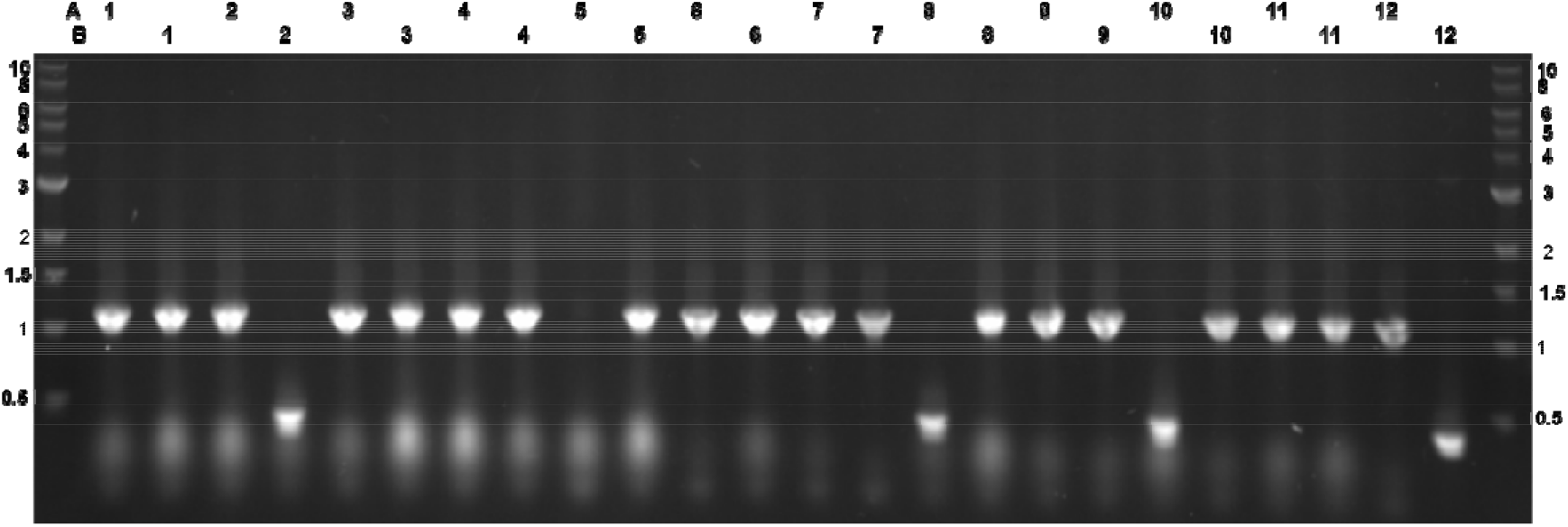
Agarose gel of colony PCR outcome for pNIC-NHSTIIT-sfGFP clones. Wells A1-B11 are pNIC-NHSTIIT-sfGFP clones and B12 is a pNIC-NHSTIIT-empty vector control. In total 19 of the 23 pNIC-NHSTIIT-sfGFP clones can be seen to contain the expected size insert of 1.1 kb.

**Supplementary Figure 2.**
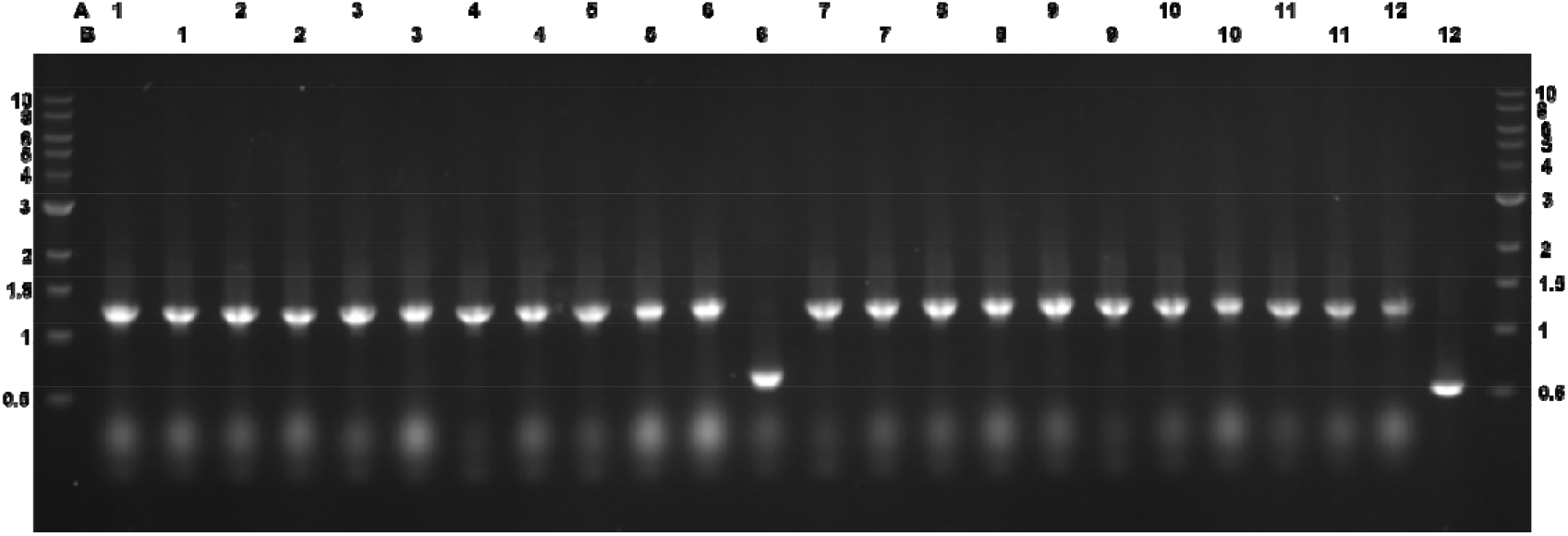
Agarose gel of colony PCR outcome for pNIC-HIS6-SUMO-sfGFP clones. Wells A1-B11 are pNIC-6HIS-SUMO-sfGFP clones and B12 is a pNIC-6HIS-SUMO-empty vector control. In total 22 of the 23 pNIC-6HIS-SUMO-sfGFP clones can be seen to contain the expected size insert of 1.3 kb.

**Supplementary Figure 3.**
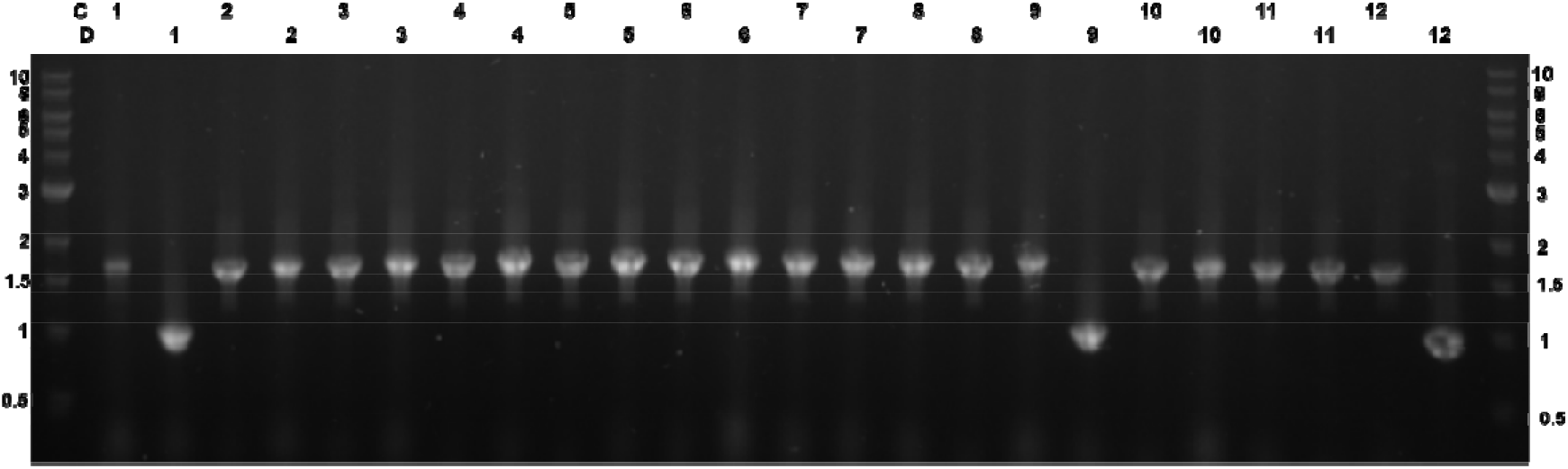
Agarose gel of colony PCR outcome for pNIC-HIS6-GST-TEV-sfGFP clones. Wells C1-D11 are pNIC-6HIS-GST-TEV-sfGFP clones and D12 is a pNIC-6HIS-GST-TEV-empty vector control. In total 21 of the 23 pNIC-6HIS-SUMO-sfGFP clones can be seen to contain the expected size insert of 1.7 kb.

**Supplementary Figure 4.**
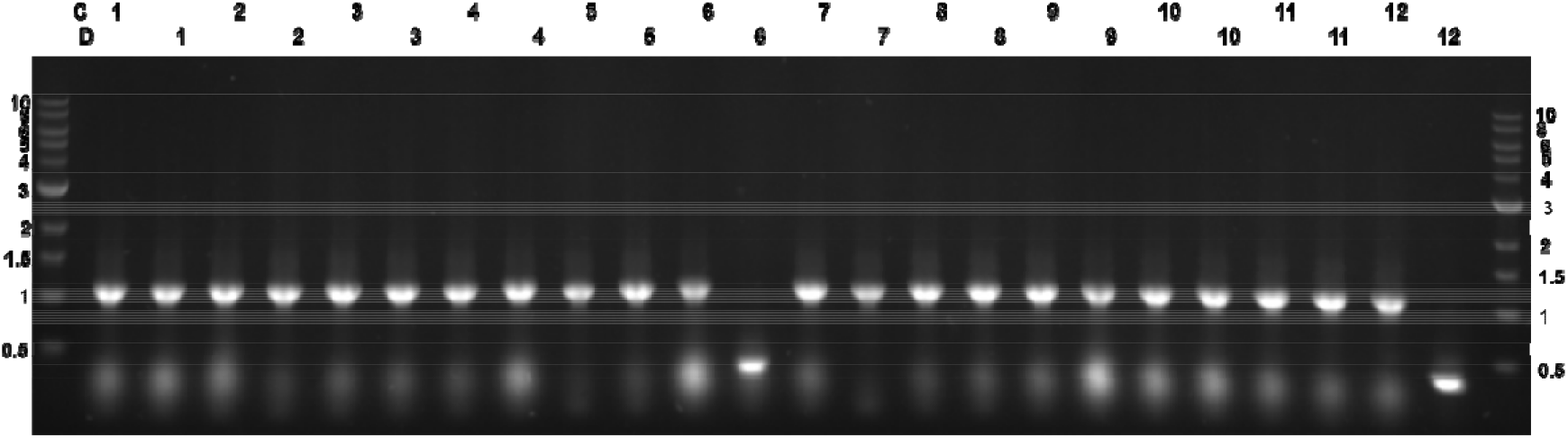
Agarose gel of colony PCR outcome for pNIC-sfGFP-CTH10STII clones. Wells C1-D11 are pNIC-sfGFP-CTH10STII clones and B12 is a pNIC-CTH10STII-empty vector control. In total 22 of the 23 pNIC-sfGFP-CTH10STII clones can be seen to contain the expected size insert of 1.1 kb.

**Supplementary Figure 5.**
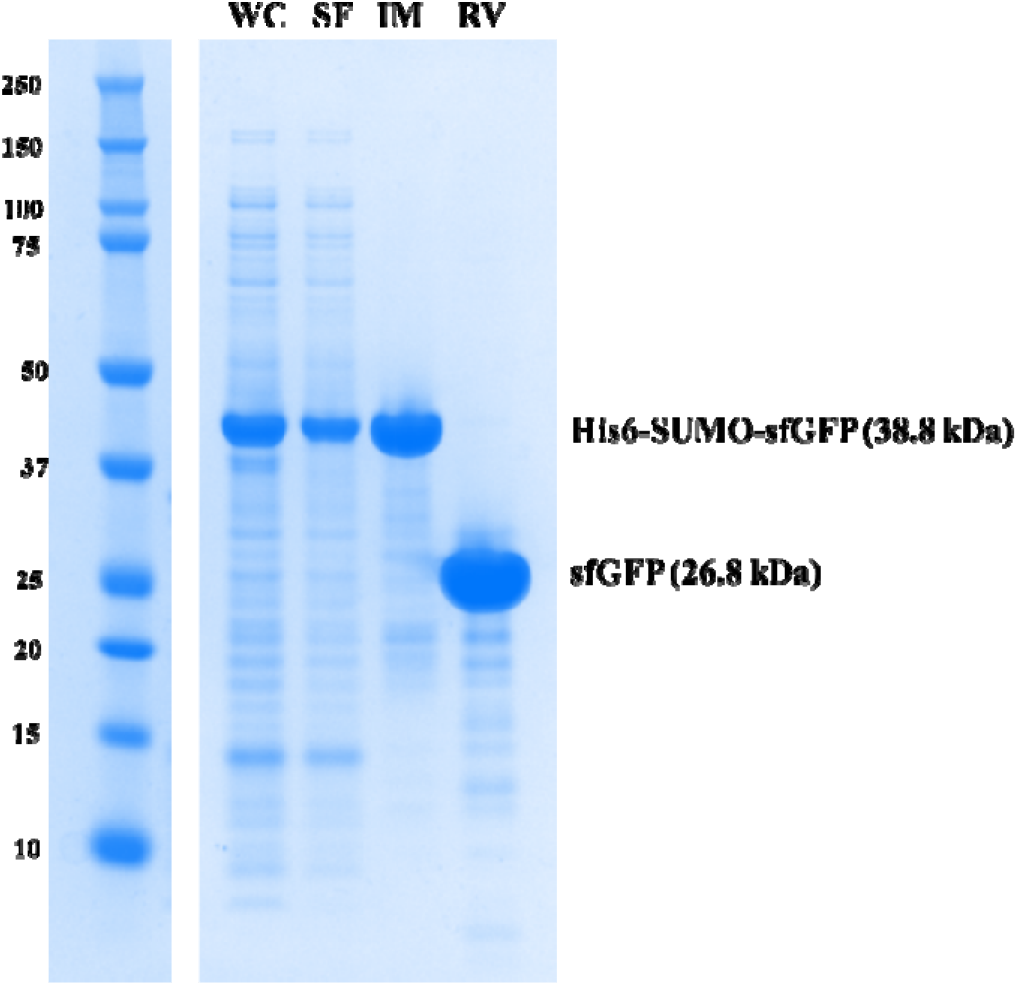
SDS-PAGE of the pNIC-His6-SUMO-sfGFP construct protein purification, where WC = whole cell lysate, SF = soluble fraction obtained after centrifugation, IM = 500 mM imidazole His Gravitrap eluate, RV = His Gravitrap flow-through after tag removal by protease cleavage.

**Supplementary Figure 6.**
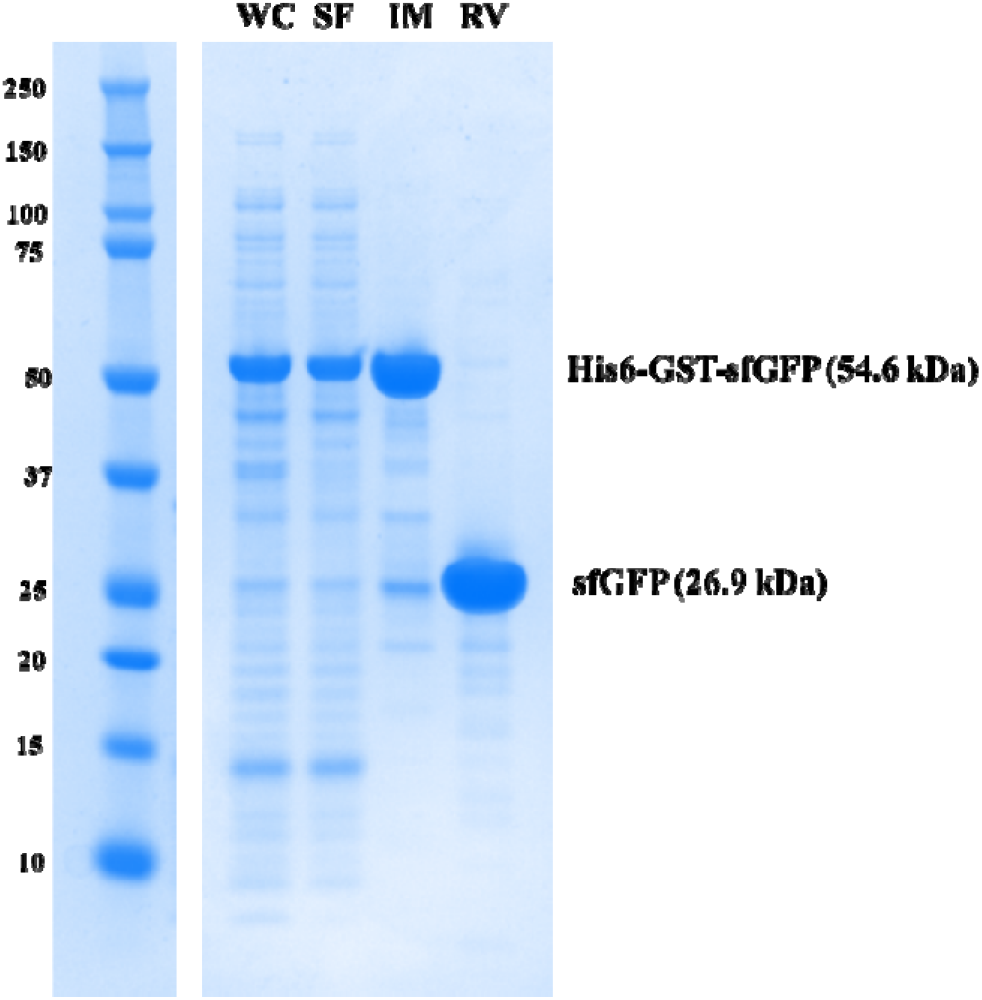
SDS-PAGE of the pNIC-His6-GST-TEV-sfGFP construct protein purification, where WC = whole cell lysate, SF = soluble fraction obtained after centrifugation, IM = 500 mM imidazole His Gravitrap eluate, RV = His Gravitrap flow-through after tag removal by protease cleavage.

**Supplementary Figure 7.**
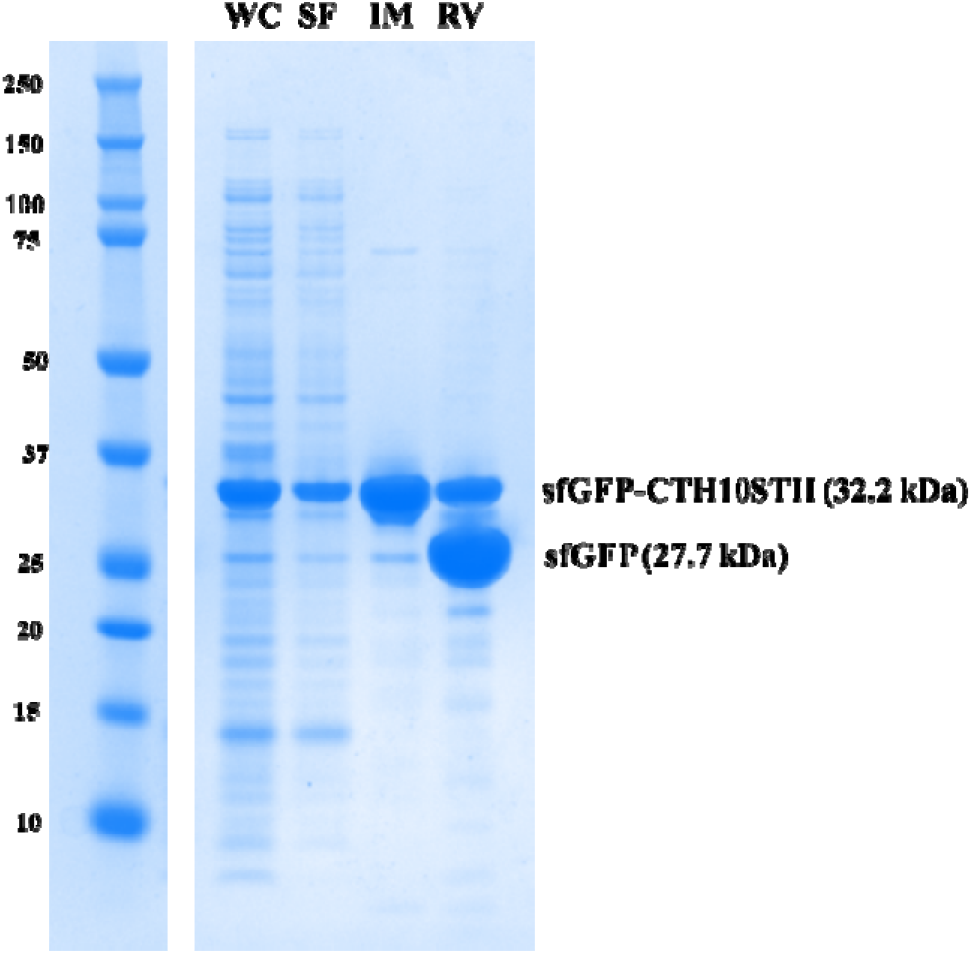
SDS-PAGE of the pNIC-CTH10STII-sfGFP construct protein purification, where WC = whole cell lysate, SF = soluble fraction obtained after centrifugation, IM = 500 mM imidazole His Gravitrap eluate, RV = His Gravitrap flow-through after tag removal by protease cleavage.

