## Supplementary material for "A selection of Golden Gate vectors to simplify recombinant protein production in *Escherichia coli*": Manuscript Figures

### Slide 1
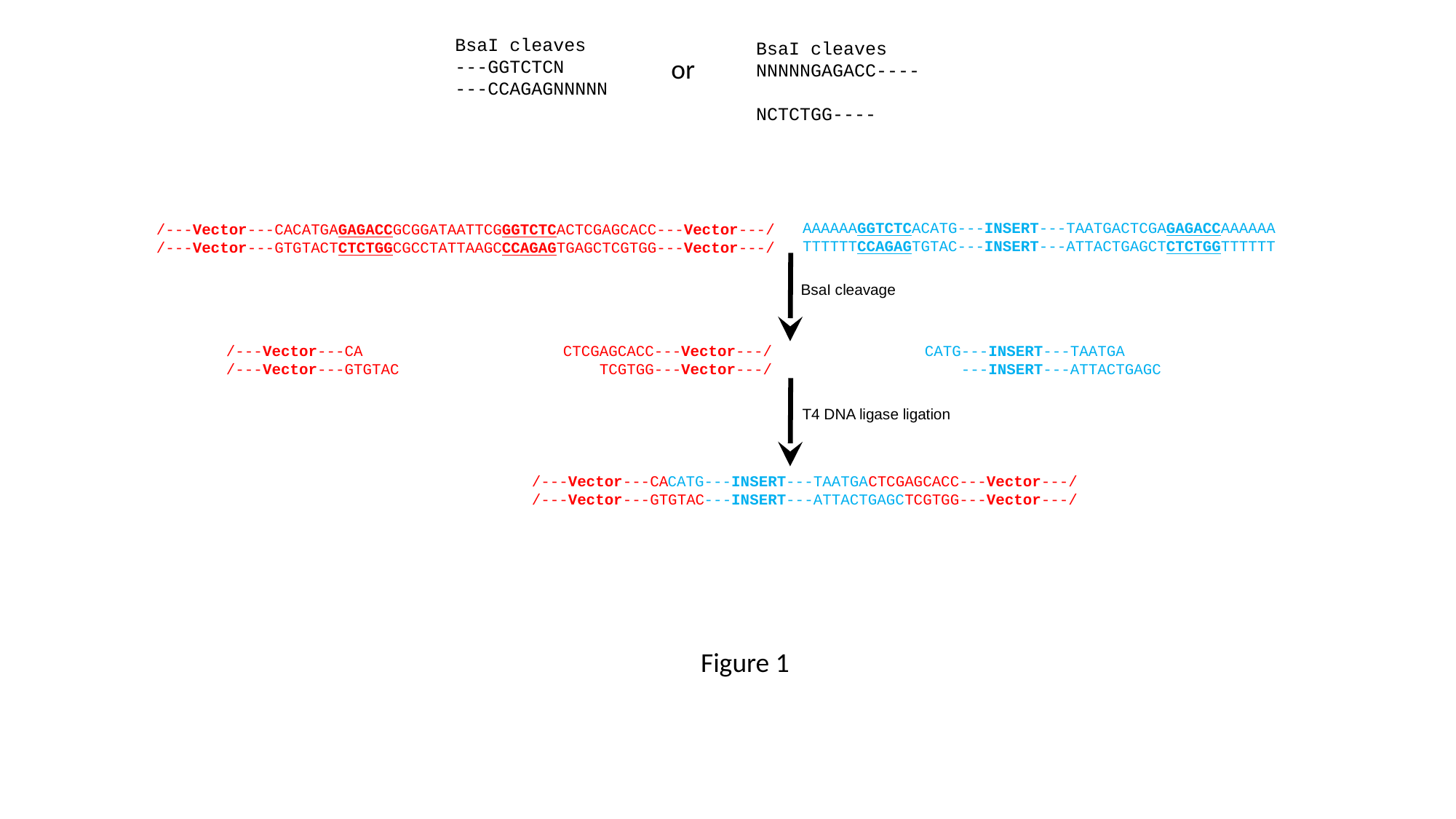

BsaI cleaves
---GGTCTCN
---CCAGAGNNNNN
BsaI cleaves
NNNNNGAGACC----
 NCTCTGG----
or
AAAAAAGGTCTCACATG---INSERT---TAATGACTCGAGAGACCAAAAAA
TTTTTTCCAGAGTGTAC---INSERT---ATTACTGAGCTCTCTGGTTTTTT
/---Vector---CACATGAGAGACCGCGGATAATTCGGGTCTCACTCGAGCACC---Vector---/
/---Vector---GTGTACTCTCTGGCGCCTATTAAGCCCAGAGTGAGCTCGTGG---Vector---/
BsaI cleavage
CATG---INSERT---TAATGA
 ---INSERT---ATTACTGAGC
/---Vector---CA CTCGAGCACC---Vector---/
/---Vector---GTGTAC TCGTGG---Vector---/
T4 DNA ligase ligation
/---Vector---CA CTCGAGCACC---Vector---/
/---Vector---GTGTAC TCGTGG---Vector---/
CATG---INSERT---TAATGA
 ---INSERT---ATTACTGAGC
Figure 1

### Slide 2
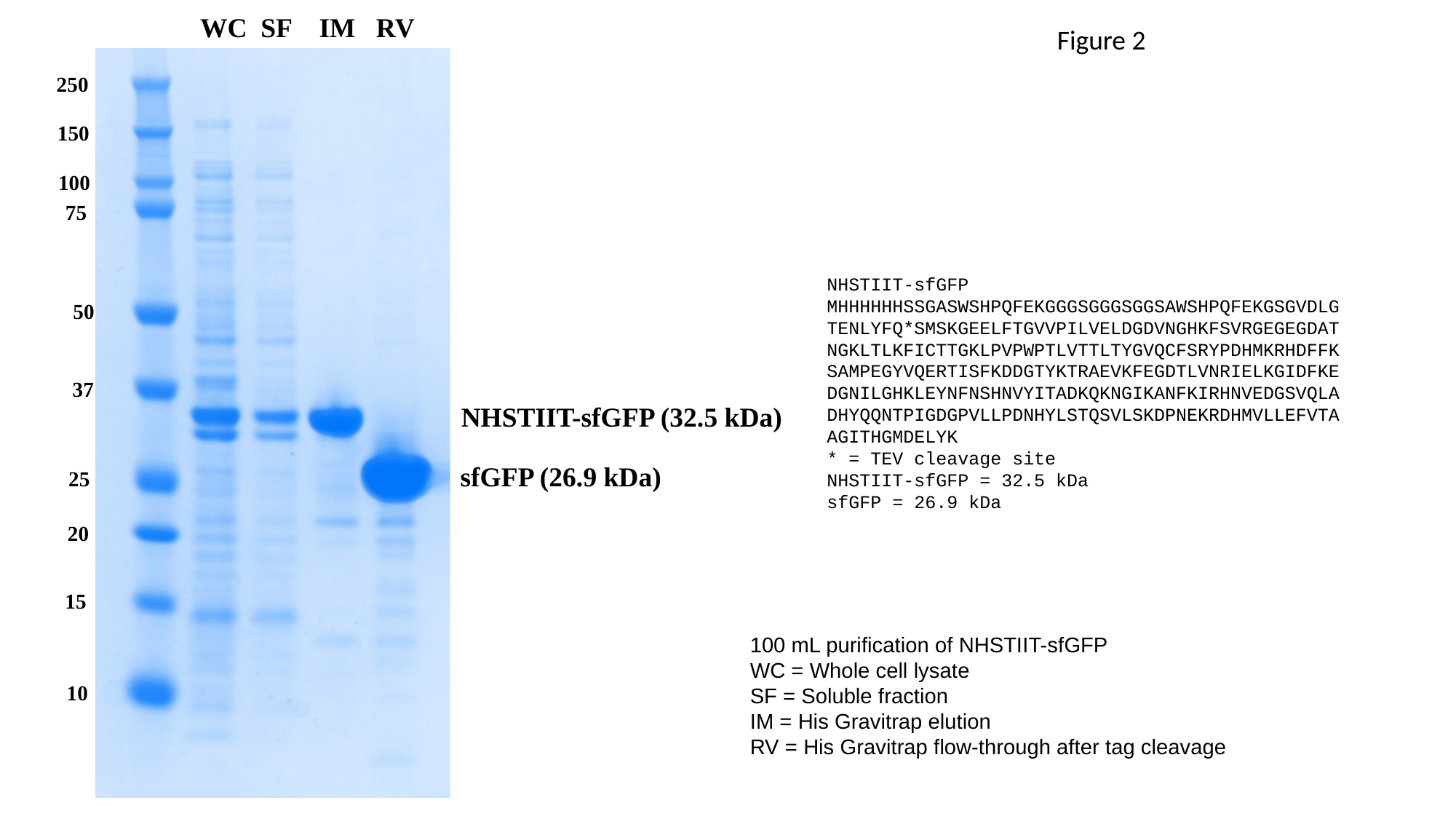

WC SF IM RV
Figure 2
250
150
100
75
NHSTIIT-sfGFP
MHHHHHHSSGASWSHPQFEKGGGSGGGSGGSAWSHPQFEKGSGVDLGTENLYFQ*SMSKGEELFTGVVPILVELDGDVNGHKFSVRGEGEGDATNGKLTLKFICTTGKLPVPWPTLVTTLTYGVQCFSRYPDHMKRHDFFKSAMPEGYVQERTISFKDDGTYKTRAEVKFEGDTLVNRIELKGIDFKEDGNILGHKLEYNFNSHNVYITADKQKNGIKANFKIRHNVEDGSVQLADHYQQNTPIGDGPVLLPDNHYLSTQSVLSKDPNEKRDHMVLLEFVTAAGITHGMDELYK
* = TEV cleavage site
NHSTIIT-sfGFP = 32.5 kDa
sfGFP = 26.9 kDa
50
37
NHSTIIT-sfGFP (32.5 kDa)
sfGFP (26.9 kDa)
25
20
15
100 mL purification of NHSTIIT-sfGFP
WC = Whole cell lysate
SF = Soluble fraction
IM = His Gravitrap elution
RV = His Gravitrap flow-through after tag cleavage
10

### Slide 3
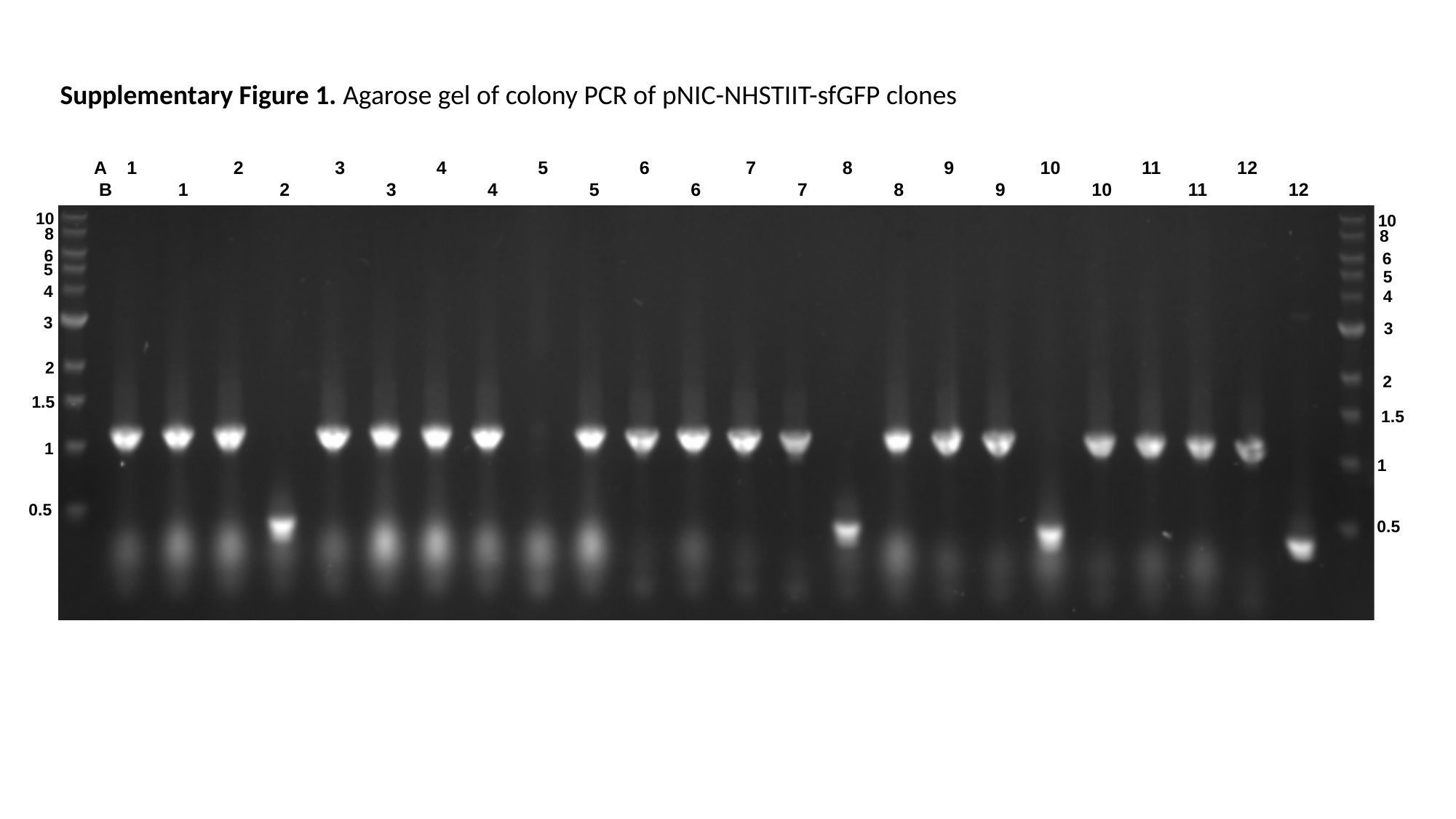

Supplementary Figure 1. Agarose gel of colony PCR of pNIC-NHSTIIT-sfGFP clones
A 1 2 3 4 5 6 7 8 9 10 11 12
 B 1 2 3 4 5 6 7 8 9 10 11 12
10
10
8
8
6
6
5
5
4
4
3
3
2
2
1.5
1.5
1
1
0.5
0.5

### Slide 4
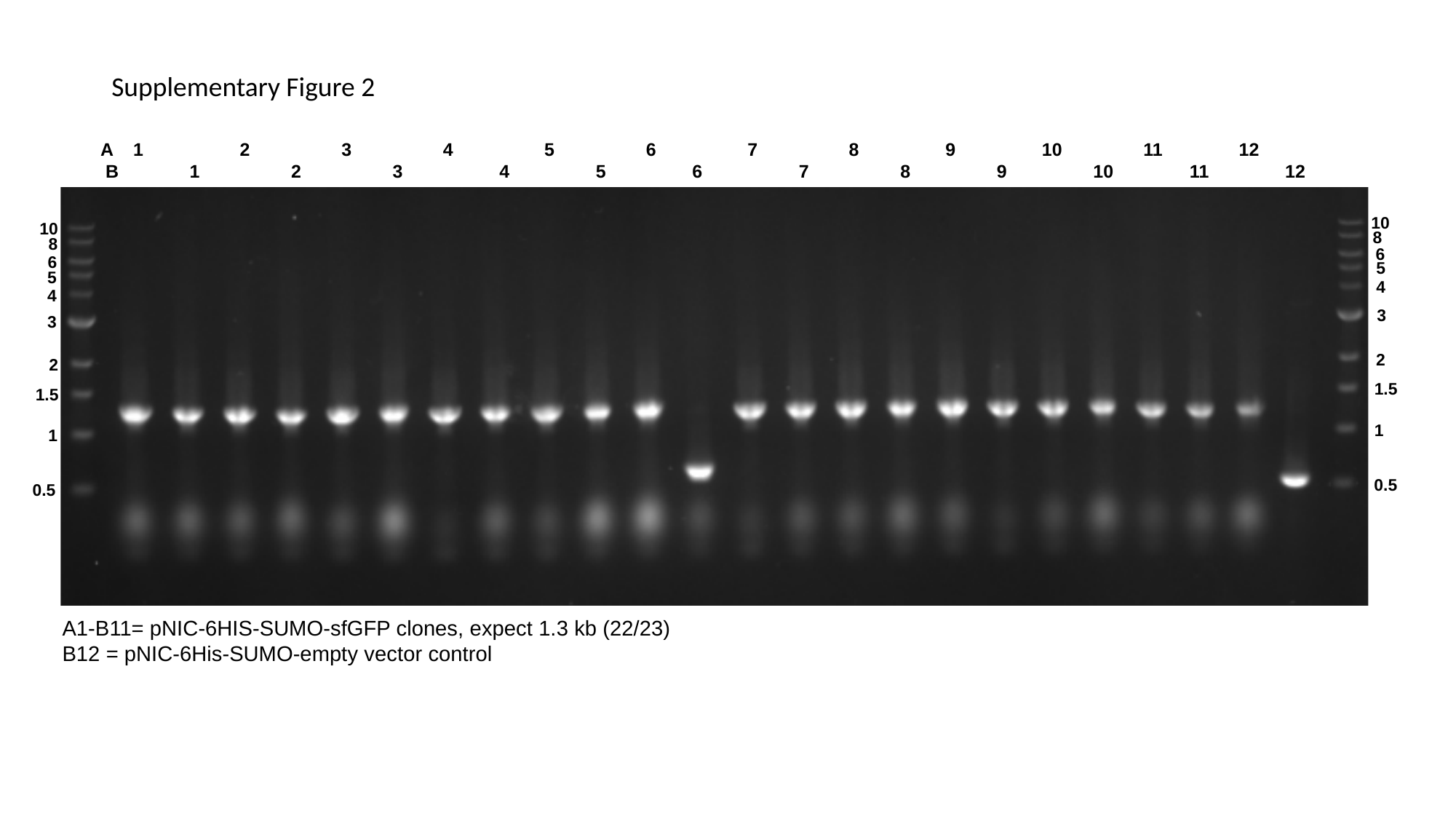

Supplementary Figure 2
A 1 2 3 4 5 6 7 8 9 10 11 12
 B 1 2 3 4 5 6 7 8 9 10 11 12
10
10
8
8
6
6
5
5
4
4
3
3
2
2
1.5
1.5
1
1
0.5
0.5
A1-B11= pNIC-6HIS-SUMO-sfGFP clones, expect 1.3 kb (22/23)
B12 = pNIC-6His-SUMO-empty vector control

### Slide 5
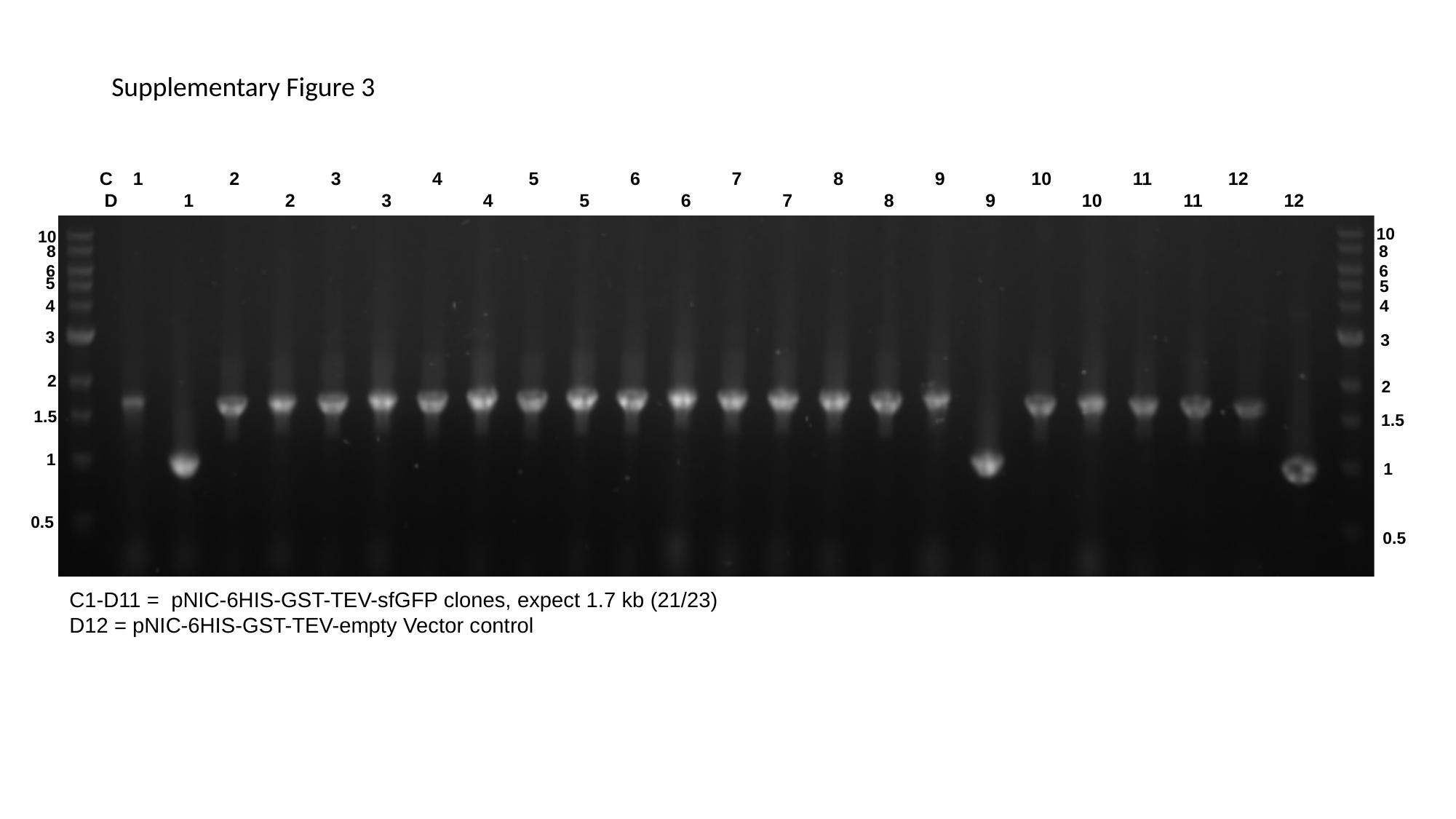

Supplementary Figure 3
C 1 2 3 4 5 6 7 8 9 10 11 12
 D 1 2 3 4 5 6 7 8 9 10 11 12
10
10
8
8
6
6
5
5
4
4
3
3
2
2
1.5
1.5
1
1
0.5
0.5
C1-D11 = pNIC-6HIS-GST-TEV-sfGFP clones, expect 1.7 kb (21/23)
D12 = pNIC-6HIS-GST-TEV-empty Vector control

### Slide 6
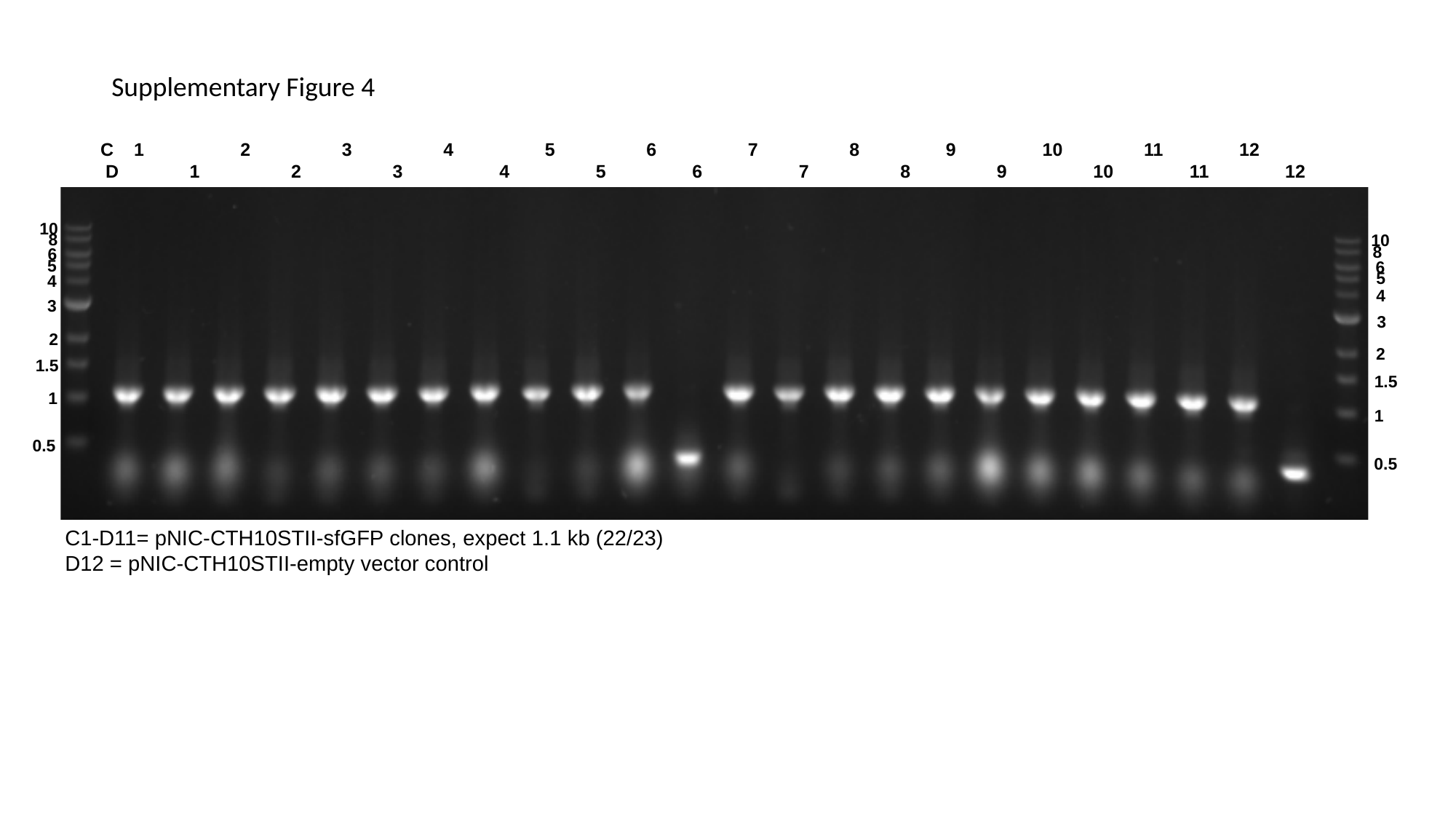

Supplementary Figure 4
C 1 2 3 4 5 6 7 8 9 10 11 12
 D 1 2 3 4 5 6 7 8 9 10 11 12
10
8
10
8
6
5
6
5
4
4
3
3
2
2
1.5
1.5
1
1
0.5
0.5
C1-D11= pNIC-CTH10STII-sfGFP clones, expect 1.1 kb (22/23)
D12 = pNIC-CTH10STII-empty vector control

### Slide 7
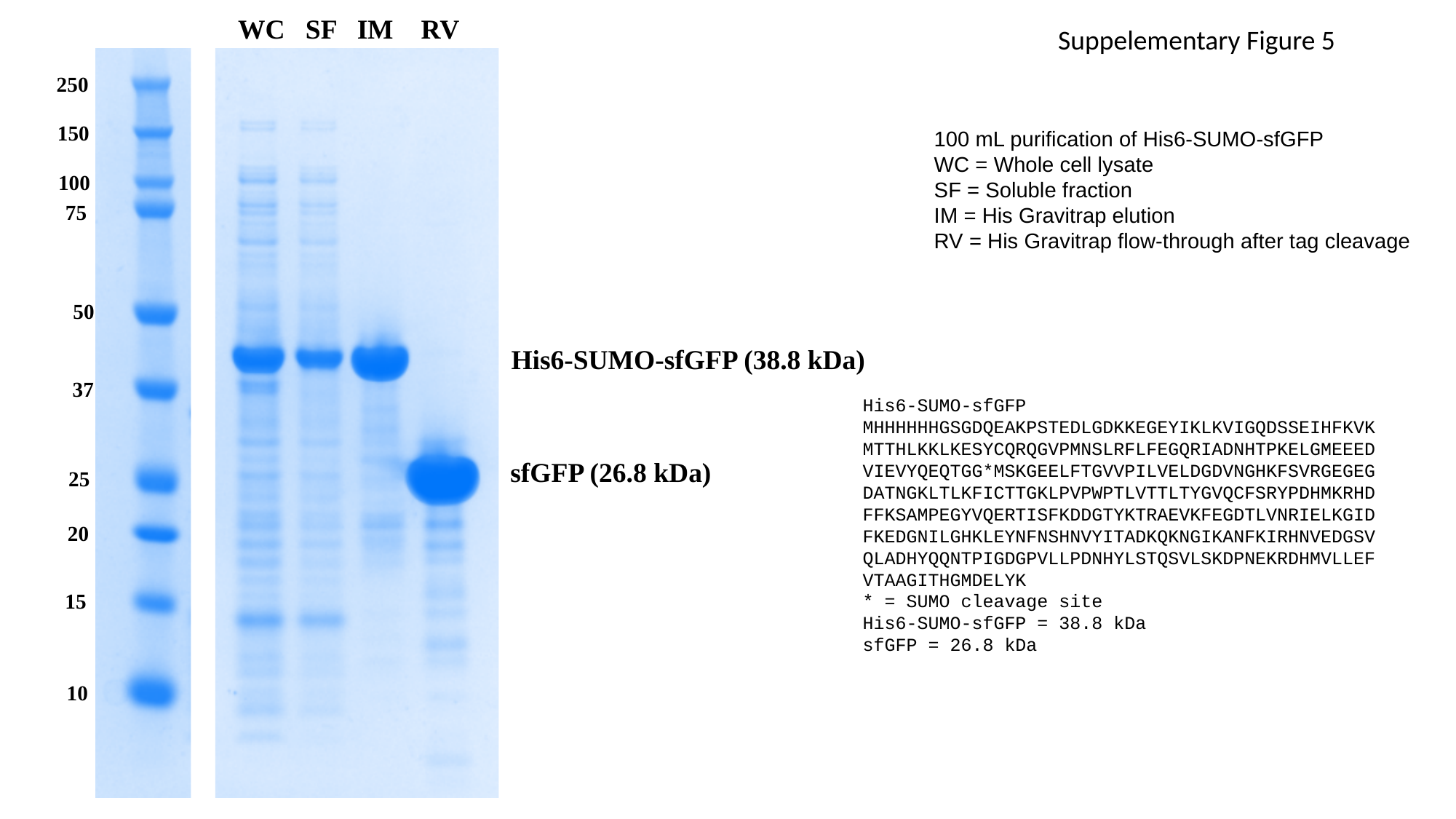

WC SF IM RV
Suppelementary Figure 5
250
150
100 mL purification of His6-SUMO-sfGFP
WC = Whole cell lysate
SF = Soluble fraction
IM = His Gravitrap elution
RV = His Gravitrap flow-through after tag cleavage
100
75
50
His6-SUMO-sfGFP (38.8 kDa)
37
His6-SUMO-sfGFP
MHHHHHHGSGDQEAKPSTEDLGDKKEGEYIKLKVIGQDSSEIHFKVKMTTHLKKLKESYCQRQGVPMNSLRFLFEGQRIADNHTPKELGMEEEDVIEVYQEQTGG*MSKGEELFTGVVPILVELDGDVNGHKFSVRGEGEGDATNGKLTLKFICTTGKLPVPWPTLVTTLTYGVQCFSRYPDHMKRHDFFKSAMPEGYVQERTISFKDDGTYKTRAEVKFEGDTLVNRIELKGIDFKEDGNILGHKLEYNFNSHNVYITADKQKNGIKANFKIRHNVEDGSVQLADHYQQNTPIGDGPVLLPDNHYLSTQSVLSKDPNEKRDHMVLLEFVTAAGITHGMDELYK
* = SUMO cleavage site
His6-SUMO-sfGFP = 38.8 kDa
sfGFP = 26.8 kDa
sfGFP (26.8 kDa)
25
20
15
10

### Slide 8
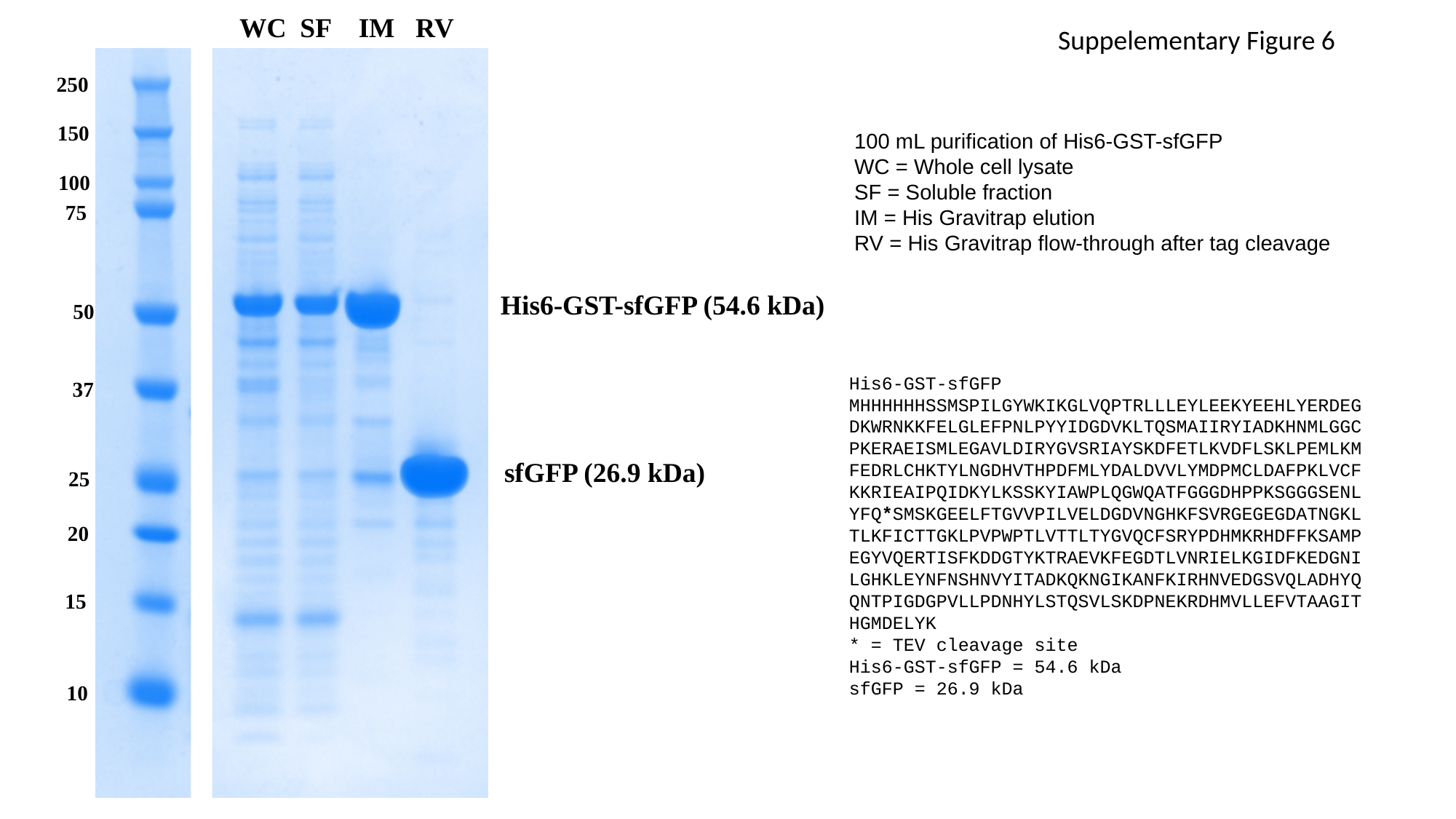

WC SF IM RV
Suppelementary Figure 6
250
150
100 mL purification of His6-GST-sfGFP
WC = Whole cell lysate
SF = Soluble fraction
IM = His Gravitrap elution
RV = His Gravitrap flow-through after tag cleavage
100
75
His6-GST-sfGFP (54.6 kDa)
50
His6-GST-sfGFP
MHHHHHHSSMSPILGYWKIKGLVQPTRLLLEYLEEKYEEHLYERDEGDKWRNKKFELGLEFPNLPYYIDGDVKLTQSMAIIRYIADKHNMLGGCPKERAEISMLEGAVLDIRYGVSRIAYSKDFETLKVDFLSKLPEMLKMFEDRLCHKTYLNGDHVTHPDFMLYDALDVVLYMDPMCLDAFPKLVCFKKRIEAIPQIDKYLKSSKYIAWPLQGWQATFGGGDHPPKSGGGSENLYFQ*SMSKGEELFTGVVPILVELDGDVNGHKFSVRGEGEGDATNGKLTLKFICTTGKLPVPWPTLVTTLTYGVQCFSRYPDHMKRHDFFKSAMPEGYVQERTISFKDDGTYKTRAEVKFEGDTLVNRIELKGIDFKEDGNILGHKLEYNFNSHNVYITADKQKNGIKANFKIRHNVEDGSVQLADHYQQNTPIGDGPVLLPDNHYLSTQSVLSKDPNEKRDHMVLLEFVTAAGITHGMDELYK
* = TEV cleavage site
His6-GST-sfGFP = 54.6 kDa
sfGFP = 26.9 kDa
37
sfGFP (26.9 kDa)
25
20
15
10

### Slide 9
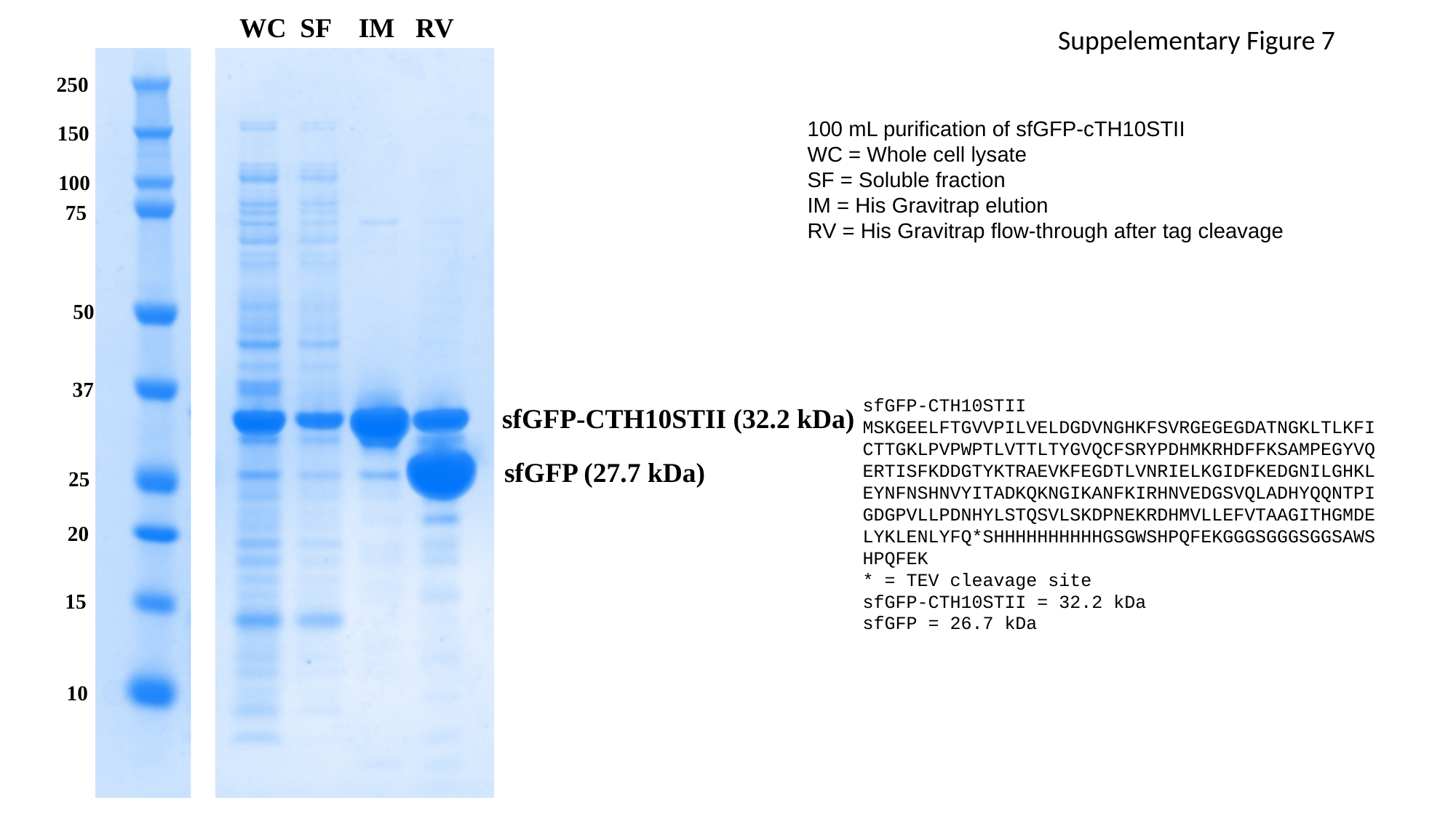

WC SF IM RV
Suppelementary Figure 7
250
100 mL purification of sfGFP-cTH10STII
WC = Whole cell lysate
SF = Soluble fraction
IM = His Gravitrap elution
RV = His Gravitrap flow-through after tag cleavage
150
100
75
50
37
sfGFP-CTH10STII
MSKGEELFTGVVPILVELDGDVNGHKFSVRGEGEGDATNGKLTLKFICTTGKLPVPWPTLVTTLTYGVQCFSRYPDHMKRHDFFKSAMPEGYVQERTISFKDDGTYKTRAEVKFEGDTLVNRIELKGIDFKEDGNILGHKLEYNFNSHNVYITADKQKNGIKANFKIRHNVEDGSVQLADHYQQNTPIGDGPVLLPDNHYLSTQSVLSKDPNEKRDHMVLLEFVTAAGITHGMDELYKLENLYFQ*SHHHHHHHHHHGSGWSHPQFEKGGGSGGGSGGSAWSHPQFEK
* = TEV cleavage site
sfGFP-CTH10STII = 32.2 kDa
sfGFP = 26.7 kDa
sfGFP-CTH10STII (32.2 kDa)
sfGFP (27.7 kDa)
25
20
15
10

### Slide 10
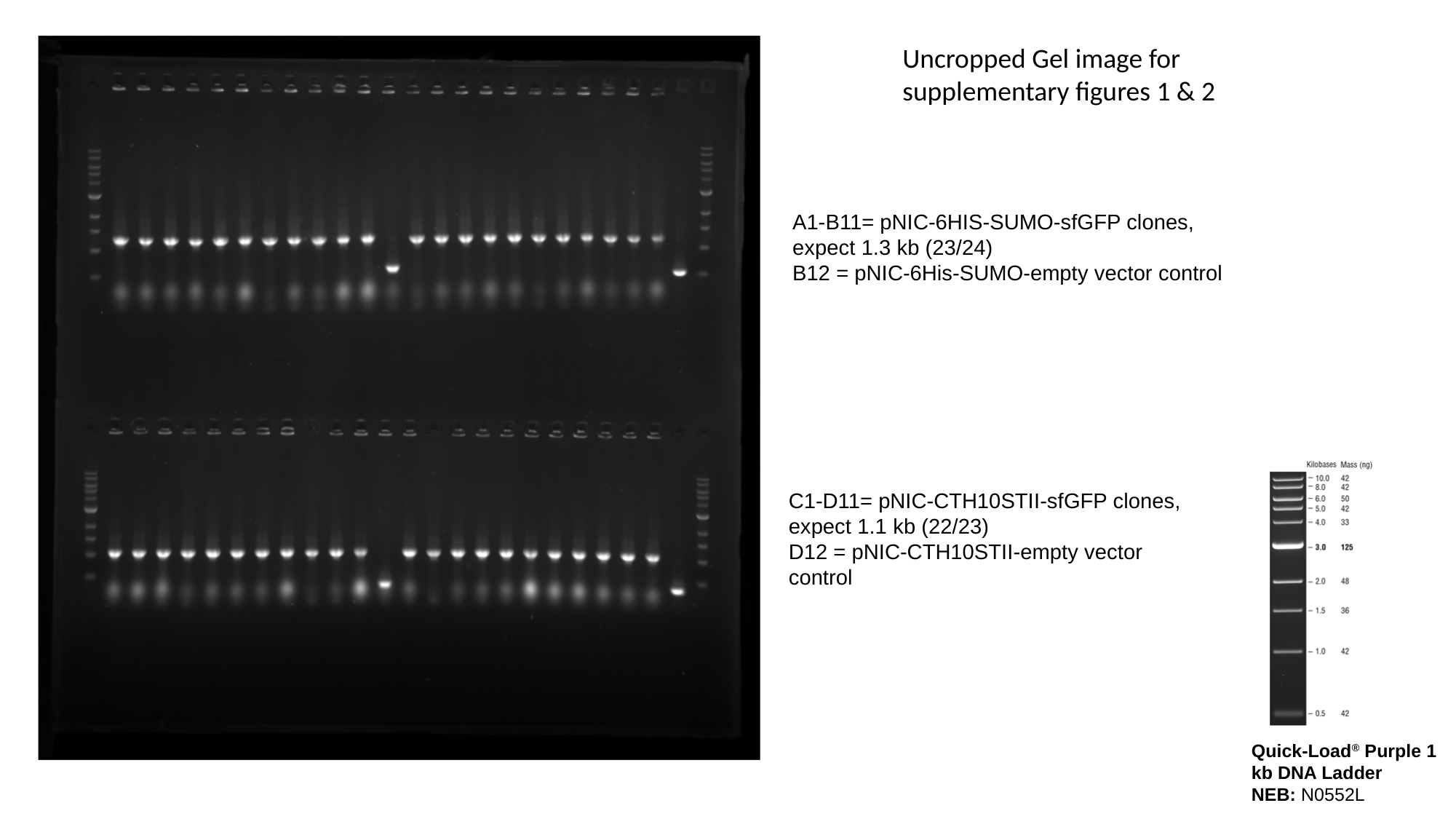

Uncropped Gel image for supplementary figures 1 & 2
A1-B11= pNIC-6HIS-SUMO-sfGFP clones, expect 1.3 kb (23/24)
B12 = pNIC-6His-SUMO-empty vector control
C1-D11= pNIC-CTH10STII-sfGFP clones, expect 1.1 kb (22/23)
D12 = pNIC-CTH10STII-empty vector control
Quick-Load^®^ Purple 1 kb DNA Ladder
NEB: N0552L

### Slide 11
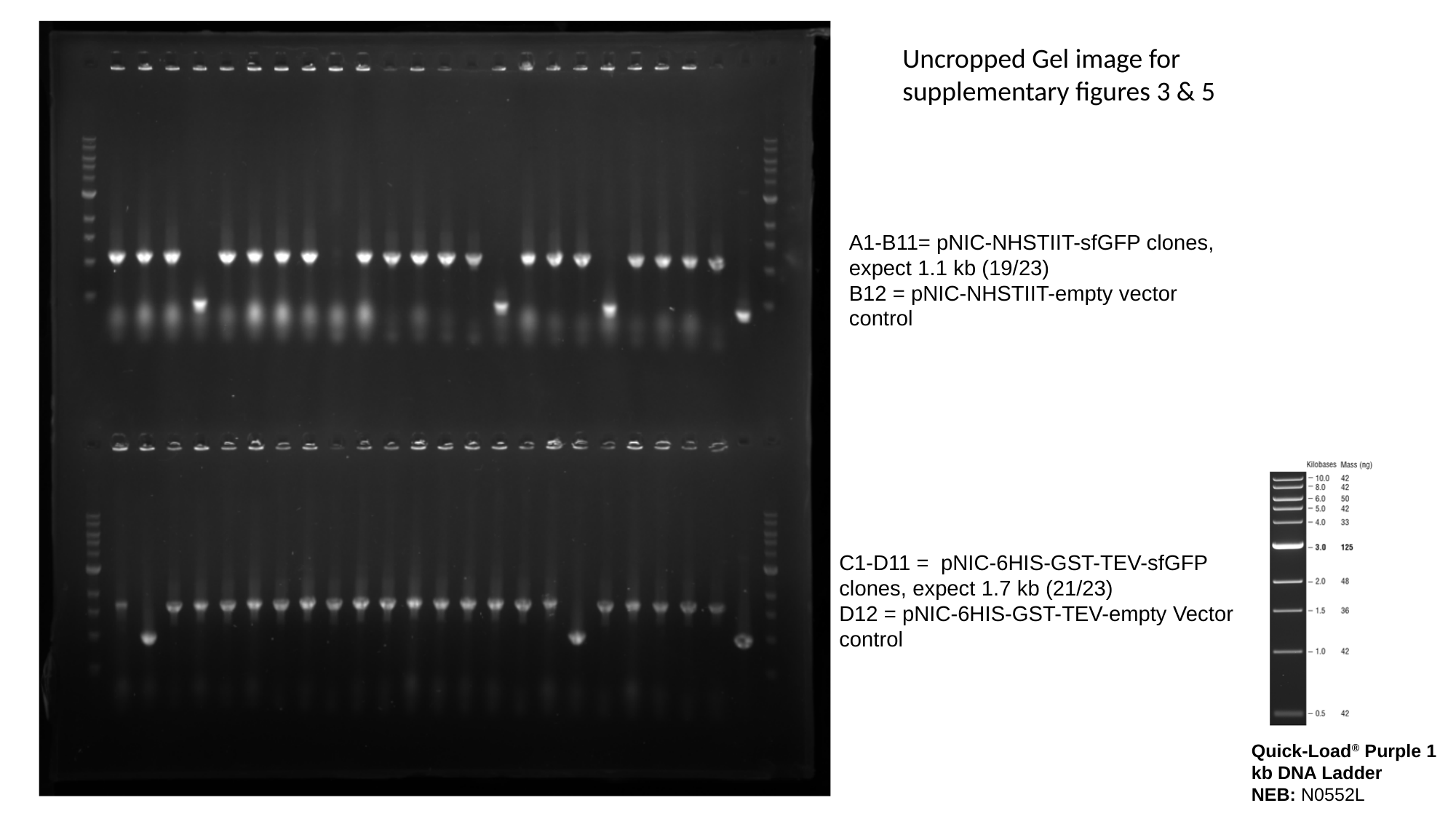

Uncropped Gel image for supplementary figures 3 & 5
A1-B11= pNIC-NHSTIIT-sfGFP clones, expect 1.1 kb (19/23)
B12 = pNIC-NHSTIIT-empty vector control
C1-D11 = pNIC-6HIS-GST-TEV-sfGFP clones, expect 1.7 kb (21/23)
D12 = pNIC-6HIS-GST-TEV-empty Vector control
Quick-Load^®^ Purple 1 kb DNA Ladder
NEB: N0552L

### Slide 12
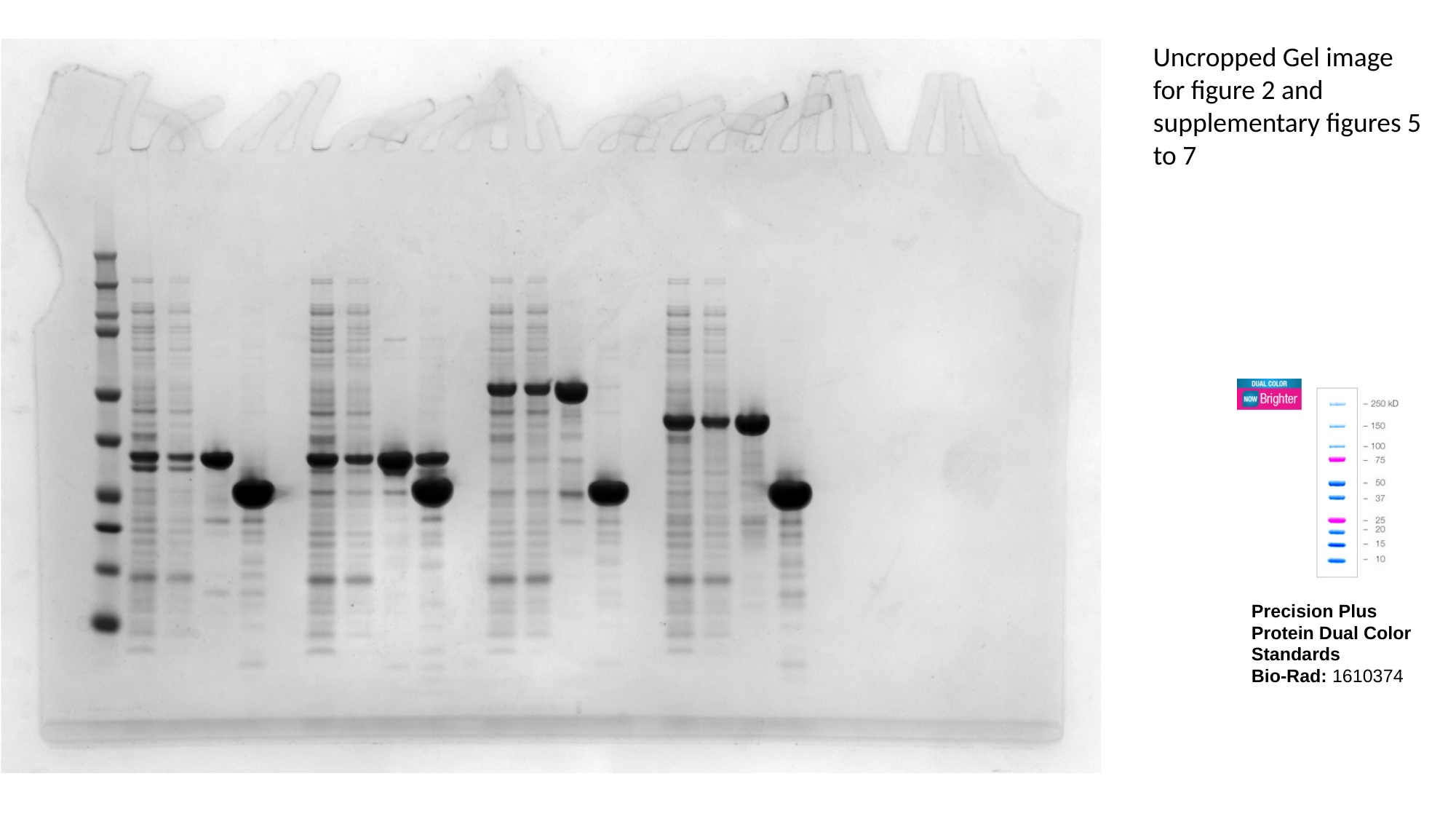

Uncropped Gel image for figure 2 and supplementary figures 5 to 7
Precision Plus Protein Dual Color Standards
Bio-Rad: 1610374
